# Multimodal single cell-resolved spatial proteomics reveals pancreatic tumor heterogeneity

**DOI:** 10.1101/2023.11.04.565590

**Authors:** Yanfen Xu, Xi Wang, Yuan Li, Yiheng Mao, Yiran Su, Yun Yang, Weina Gao, Changying Fu, Wendong Chen, Xueting Ye, Fuchao Liang, Panzhu Bai, Ying Sun, Ruilian Xu, Ruijun Tian

## Abstract

Despite the advances in antibody-guided cell typing and mass spectrometry-based proteomics, their integration is hindered by challenges for processing rare cells in the heterogeneous tissue context. Here, we introduce Spatial and Cell-type Proteomics (SCPro), which combines multiplexed imaging and flow cytometry with ion exchange-based protein aggregation capture technology to characterize spatial proteome heterogeneity with single cell resolution. The SCPro was employed to explore the pancreatic tumor microenvironment and revealed the spatial alternations of over 5,000 proteins by automatically dissecting up to 100 single cells guided by multi-color imaging of centimeter-scale formalin-fixed, paraffin-embedded tissue slide. To enhance cell-type resolution, we characterized the proteome of 14 different cell types by sorting up to 1,000 cells from the same tumor, which allows us to deconvolute the spatial distribution of immune cell subtypes and leads to the discovery of a novel subtype of regulatory T cells. Together, the SCPro provides a multimodal spatial proteomics approach for profiling tissue proteome heterogeneity.

## Main

Spatial and cell-type heterogeneity is ubiquitous in the tissue context and plays a vital role in constituting the functional diversity of human diseases^1^. While single-cell and spatial transcriptomics have made remarkable achievements in revealing the intra-tumor cell diversity, in-depth profiling of its protein basis at single-cell resolution remains challenging^1,2^. Antibody recognition-based cell typing technologies by dissociating the tissue specimens into single-cell suspension^3^ or multiplexed imaging analysis of the tissue microenvironment have been widely used to analyze dozens of proteins at the cellular or even subcellular resolution^4,5^. However, these targeted methods are limited by the number of available antibodies, thus falling short of comprehensively capturing the intricate cellular proteome. Recently, tremendous progress in mass spectrometry (MS)-based proteomics has rendered it a powerful tool for exploring the proteome in unbiased and global manners^6^.

Spatially resolved proteomics based on various microdissection techniques has made tremendous progress in profiling of thousands of proteins while preserving spatial information^7–10^. Guided by hematoxylin-eosin (H&E)- or immunohistochemical (IHC)-stained images, laser microdissection (LMD)-based spatial proteomics has been successfully demonstrated in providing unprecedented insights into the heterogeneous tissue context of lethal disease tissue samples with cell-type resolution. Starting with either fresh frozen or formalin-fixed, paraffin-embedded (FFPE) tissue slice samples, LMD-based spatial proteomics has been widely applied to ovarian cancer^11^, colon cancer^12,13^, tuberculosis^14^, pancreatic cancer^15^, COVID-19^16,17^, etc. However, the spatial resolution of these studies largely depends on experienced pathological examination, subjective cell typing, and time-consuming manual selection of tissue regions of interest (ROI). Furthermore, the large-scale ROI pieces obtained from these studies commonly have low cell-type resolution, which results in an averaging effect and ultimately blurs the spatial and cell-type information.

Efforts have been made to improve the precision and throughput of LMD-based spatial proteomics. Mund et al. recently reported the development of Deep Visual Proteomics (DVP) which combines artificial-intelligence (AI)-driven image-based cell segmentation and automated LMD for spatial proteomic profiling of IHC-stained melanoma tissue with single cell resolution^18^. Very recently, the DVP platform was also used to study the proteome of immunofluorescence-stained melanoma cells within the epidermal and dermal compartments of primary cutaneous melanoma^19^. Despite these pioneered advancements made in image-guided spatial proteomics, centimeter-scale multi-color IHC (mIHC) image navigation is need to precisely target diverse single cell types in the tumor microenvironment with high heterogeneity. Technical challenges, such as precise cell typing based on multiplexed imaging, navigation transfer between high-quality imaging with the coverslip and low-quality imaging with LMD microscope after removing the coverslip, and the sophisticated targeting, collecting, and processing of rare stained tissue cells (e.g., <100 cells to single cells) for spatial proteomic analysis, remain to be addressed.

Here, we introduce the Spatial and Cell-type Proteomics platform, termed SCPro (**Fig. 1**), which enables the integration of image-guided spatial proteomics and flow cytometry-based cell-type proteomics to uncover cell-type heterogeneity in tissue context. The spatial proteomics aspect of the SCPro coordinates accurately defined single cell contours of centimeter-scale mIHC images based on nuclei and cell membrane identification algorithms, automated LMD at single cell resolution with no-failure capture, ion exchange-based protein aggregation capture (iPAC) technology for integrated proteomics sample preparation and highly sensitive proteomics profiling of rare stained FFPE tissue cells (**Fig. 1a,c**). Furthermore, inspired by deconvolution algorithms that align cell-type information in spatial transcriptomics using single-cell RNA sequencing data as a reference^20,21^, we seek to extend the cell-type resolution of the SCPro by incorporating flow cytometry-based proteomics data of 14 distinct cell types as a reference map to deconvolute the cell type composition and proportion in spatial proteome profiles (**Fig. 1b-d**). Collectively, we applied the SCPro platform to explore the spatial proteome heterogeneity of mouse pancreatic tumor microenvironment (TME) and identified a novel subtype of regulatory T cells.

**Fig. 1.**
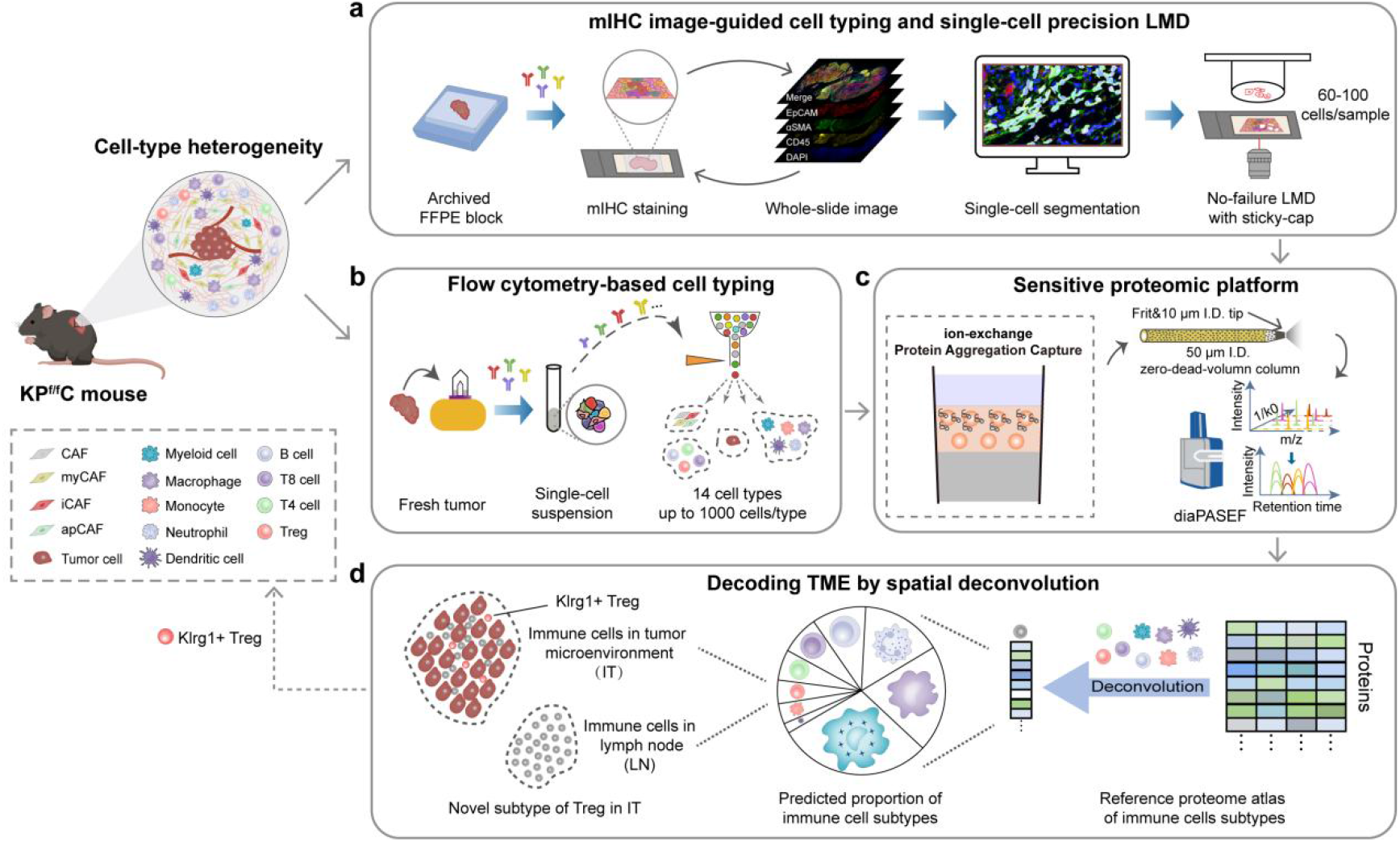
Concept and workflow of the SCPro platform. The SCPro platform integrates multiple modules. **a**, Antibody-guided cell typing based on high-quality multiplexed imaging and automated LMD with single-cell resolution. **b**, Flow cytometry-based cell typing. **c**, Ultra-high-sensitivity proteomics platform combining with ion exchange-based protein aggregation capture sample preparation, low-flow chromatography, and high-sensitivity mass spectrometry data acquisition. **d**, Decoding the pancreatic tumor microenvironment through spatial deconvolution. CAF, cancer-associated fibroblast; iCAF, inflammatory CAF; myCAF, myofibroblastic CAF; apCAF, antigen-presenting CAF; Treg, regulatory T cell; FFPE, formalin-fixed paraffin-embedded; mIHC, multiplexed immunohistochemical; LMD, laser microdissection; TME, tumor microenvironment.

## Results

### iPAC enables in-depth spatial proteomic profiling of <100 cells in FFPE tissue slice

One of the key technical hindrances for integrating antibody recognition-based cell typing technologies and MS-based proteomics to uncover tissue proteome heterogeneity is the manipulation of rare cells. The development of integrated proteomics sample preparation platform for processing limited number of cells is therefore crucial for avoiding sample loss and achieving highly sensitive proteome profiling^22^. Notably, in the case of image navigation-based spatial proteomics, the tissue slides are often FFPE-processed and stained for visualization and cell typing. It is inevitable for hydrophobic chemical dyes and surfactants to be introduced into the tissue lysates as contaminants, which cannot be effectively removed even with the peptide clean-up steps. The accumulation of these contaminants compromises chromatographic performance and significantly influences the high-sensitive MS analysis over time^13^. Here, we introduce the solid-phase extraction (SPE)-based iPAC technology for handling low quantities of cells (e.g., <100 cells to single cells), especially for stained tissue slice samples.

The iPAC device is a spintip packed with strong anion exchange (SAX) and C18 disks in tandem which stems from our previously developed fully integrated SPE-based integrated proteomics sample preparation technology, SISPROT^22,23^. Notably, we incorporated several key technical advancements to enhance the robustness of the iPAC (**Fig. 2a**). Firstly, we introduced the “carrier” surfactant N-dodecyl-β-D-maltoside (DDM) to prevent nonspecific adsorption of low nanogram-level proteins during sample processing in the iPAC spintip^24^. Secondly, we employed ion-exchange disks instead of loosely packed beads that significantly improves protein capturing efficiency. Thirdly, we implemented in-situ protein aggregation capture^25^ by simply introducing an incubation step with pure ACN after protein capture and concentration onto the SAX disks at basic pH, which well induces precipitation of proteins and facilitates extended wash to remove contaminants and pH exchange for enzymatic digestion. The resulting peptides were then salt-eluted onto C18 disks at acidic pH for desalting and ultimately transferred to a glass insert for direct injection with negligible sample loss (**Extended Data** Fig. 1a,b). Last but not least, we significantly improved the sensitivity of the LC-MS system by integrating a homemade zero-dead-volume (ZDV) 50 μm I.D. column which has a short frit made at the end of the emitter tip with neglectable dead volume and significantly improved ionization efficiency (**Fig. 2a**)^26^. The homemade ZDV column running at 100 nL/min, in turn, enables the identification of over 3,000 protein groups from 1 ng pre-digested HeLa cell samples in ddaPASEF acquisition mode, without using the match between run (MBR) algorithm (**Extended Data** Fig. 1c).

**Fig. 2.**
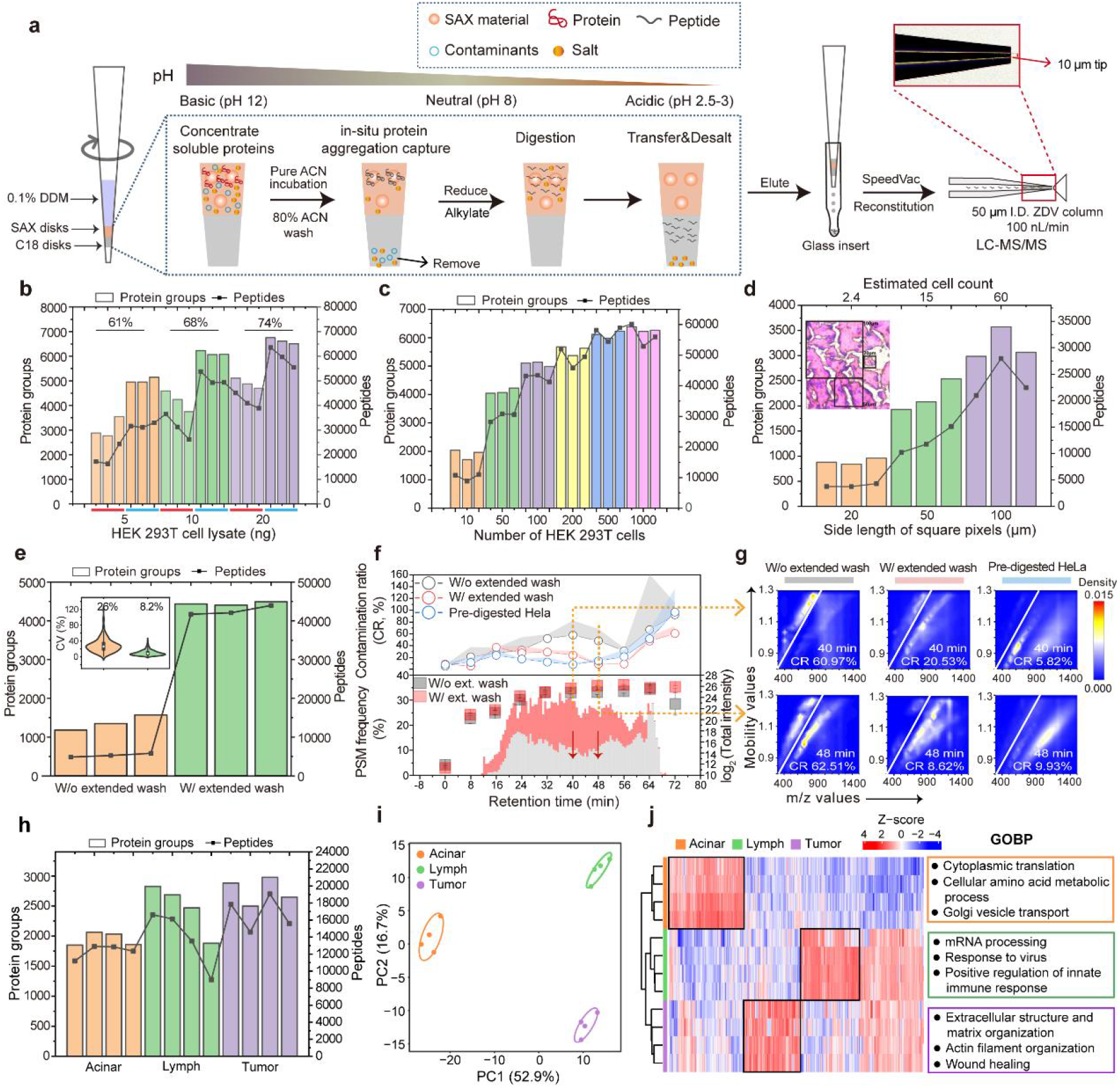
Development of the iPAC technology for processing rare stained tissue cells. **a**, Workflow and principle of iPAC technology. **b**, Identified protein groups and unique peptides from processing nanogram HEK 293T cell lysate or direct injection of an equal number of pre-digested HEK 293T peptides. 4/5 peptides were injected for iPAC processed samples. The proportion shown on the graph indicates the protein recovery rate for each group, which is determined by the ratio between the protein groups identified by the iPAC processed samples and directly injecting an equal number of peptides. **c**, Identified protein groups and unique peptides from 10 to 1000 flow cytometry-sorted HEK 293T cells. **d**, Identified protein groups and unique peptides from 20, 50, and 100 μm-side length square tissue samples of 12 μm-thick H&E-stained mouse brain. **e**, Identified protein groups and unique peptides from 12 μm-thick, 200 μm-side length square H&E-stained mouse brain samples without (W/o) or with (W/) extended wash. The insert diagram shows Coefficient of Variation (CV) distributions. **f**, Upper panel, Contamination Ratios (CR) of H&E-stained mouse brain samples without (W/o) or with (W/) extended wash along the LC gradient. The commercially available pre-digested HeLa was served as a control. The transparent shades beyond each dot-line indicate the half standard deviation of CR within each group (n = 3). Bottom panel, identified peptide-spectrum matches (PSMs) along with retention time (RT) for tissue samples without (W/o) or with (W/) extended wash. The total ion chromatogram (TIC) intensity of precursors extracted for generating heatmaps is also displayed (n = 3). **g**, Representing heatmap of precursors in the trapped ion mobility (IM)-m/z space at specific RT from one of the three replicates in each group. **h**, Identified protein groups and peptides from different regions of the KP^f/f^C mouse tissue section. **i**, Principal Component Analysis (PCA) analysis. **j**, Unsupervised clustering analysis of the significant difference proteins (ANOVA, q-value < 0.05, Fold change > 1.2). Protein expression levels were Z-scored. Top enriched Gene Ontology (GO) categories analyzed with significantly differentially expressed proteins are shown. DDM, N-dodecyl-β-D-maltoside; SAX, strong anion exchange; ZDV, zero-dead-volume.

We first evaluated the performance of the iPAC by side-by-side comparing the protein recovery rate by either processing 5 ng, 10 ng, and 20 ng of HEK 293T cell lysate or directly injecting the same amounts of pre-digested HEK 293T samples (**Fig. 2b**). More than 3,000, 4,000, and 5,000 protein groups were identified, respectively, using ddaPASEF mode without the MBR algorithm. Comparing with direct injection of 5 ng peptides, we observed a recovery rate of over 60% in terms of protein group identification (**Fig. 2b**). Notably, a higher recovery rate was obtained by increasing the cell input. Next, we assessed the performance of iPAC using 10 to 1000 flow cytometry-sorted HEK 293T cells (**Fig. 2c**). Over 2,000 and 5,000 protein groups were identified from 10 and 100 sorted cells, respectively, while deactivating the MBR algorithm to avoid overestimation of protein identification for 10-cell samples^24^. The median coefficient of variation (CV) of maxLFQ intensities within groups was less than 10%, except for the 10-cell group with a median CV of approximately 15% (**Extended Data** Fig. 1d).

We then went on to validate the performance of the iPAC for analyzing H&E-stained mouse brain tissue slices which contains much more contaminants as compared with sorted cells. Excitingly, the iPAC identified over 800, 2,000, and 3,200 protein groups from 20, 50, and 100 μm-side length square samples, respectively, corresponding to 2.4, 15, and 60 cells in volume (**Fig. 2d**). Notably, the iPAC demonstrated high quantitative reproducibility with CV values below 15% (**Extended Data** Fig. 1e). The excellent performance of the iPAC is largely attributed to the extended 80% ACN wash of aggregated proteins captured by the SAX disks, which is only available in this double-layer SPE-based sample preparation technology. As showed in **Fig. 2e**, the original version of the iPAC for processing a 200 μm-side length square slice (approximately 240 tissue cells) without an extended 80% ACN wash only identified 1,368 protein groups with a high quantitative CV of 26%, indicating a significant drop in identification compared to equal number of cultured cell samples (**Fig. 2c**). In comparison, we identified on average 4,445 protein groups and 42,595 unique peptides after adding the extended wash, representing a three-fold increase in the number of protein groups and an eight-fold increase in the number of unique peptides (**Fig. 2e**). Moreover, the median CV decreased from 26% to 8.2%, demonstrating greatly improved quantitative reproducibility (**Fig. 2e, insert**). This result is consistent with a significantly higher number of peptide spectrum matches (PSMs) across the whole LC gradient for the iPAC-processed sample with the extended wash, although the total ion chromatography (TIC) intensity was similar between samples with (W/) and without (W/O) the extended wash (**Fig. 2f, bottom panel**).

We hypothesized that this drop in protein identification without the extended wash was primarily due to the inherent nature of stained tissues, rather than sample loss. To investigate this, we evaluated the contamination ratio (CR) which is defined by analyzing the ratio of singly charged and multiple charged precursors in the ion mobility-mass spectrometry (IM-MS) heatmap across the whole LC gradient (**Fig. 2f,g, Methods**)^27^. By incorporating the extended wash, the CR significantly decreased, especially in the middle of the LC gradient from 20 to 60 minutes (**Fig. 2f, upper panel**). For example, the tissue samples showed 3 and 7 times cleaner at 40 and 48 minutes of the LC gradient, respectively, which is close to the pre-digested Hela cell results, but the CR for the sample without extended wash is more than 60% (**Fig. 2g**). To further demonstrate the robustness of the iPAC technology in processing complex biological samples, we applied it to analyze the proteome of distinct regions in pancreatic tumor tissue sections. Pancreatic cancer, with its desmoplastic and immunosuppressive tumor microenvironment (TME) characterized by cancer associated fibroblasts (CAFs) and immune cells surrounding the tumor cells, exhibits high heterogeneity and serves as a prototypical example of solid tumors with a poor prognosis^28^. We collected acinar, lymph node, and tumor regions by dissecting 100 μm-side length square tissue (approximately 100 tissue cells) from a fresh frozen tissue slice of a transgenic mouse model KP^f/f^C (Kras^LSL-G12D/+^; Trp53^flox^; Pdx1-Cre) (**Extended Data** Fig. 2a)^29^. Around 2,000 proteins were reliably identified with good quantitative reproducibility (**Fig. 2h** and **Extended Data** Fig. 2b). PCA analysis effectively differentiated cell types within the same tissue slice (**Fig. 2i**). Furthermore, we observed the enrichment of relevant biological processes (GOBP) for distinct cell types (**Fig. 2j**). Importantly, we also applied the iPAC to spatial proteomic analysis of approximately 100 cancer cells and their adjacent stromal cells dissected out based on their cell contour and spatial proximity (**Extended Data** Fig. 3). The results showed the excellent performance of the iPAC in achieving in-depth spatial proteomic profiling and recapitulating the biological features of <100 cells in stained tissue samples.

### SCPro captures the pancreatic TME with single-cell resolution

After evaluating the sensitivity of the iPAC, we then focused on accurately defining single-cell boundaries and transferring cell typing results into LMD microscope to guide automated microdissection. Here, we benchmarked the technological development and application of the SCPro utilizing a FFPE tissue block from the KP^f/f^C mouse, which well recapitulates malignant transformation process from normal acinar cells to pancreatic intraepithelial neoplasia (PanIN) and ultimately pancreatic ductal adenocarcinoma (PDAC)^30,31^. After a rigorous pathological screening process to best represent the progression of PDAC on a single tissue section, a 4-μm-thick FFPE tissue section from a 7-week-old KP^f/f^C mouse was subjected to 4-color mIHC staining. As the open-faced frame slide without the coverslip generated poor-quality images (**Extended Data** Fig. 4), navigation transfer across microscopes with and without the coverslip is a must for precise cell segmentation. Importantly, square reference shapes for image alignment were marked onto the membrane slide by LMD (**Fig. 3a**). Then, the high-quality multiplexed whole-slide image was obtained with the coverslip on the TissueFAXS system for accurate cell segmentation. After removing the coverslip from the frame slide, the cell mask was imported and aligned with the low-resolution real-time image of the LMD microscope system. Subsequently, the annotated single cell contours were isolated along the border line of cell contours by automatic LMD and captured by a sticky-cap with no-failure for downstream proteomic analysis.

**Fig. 3.**
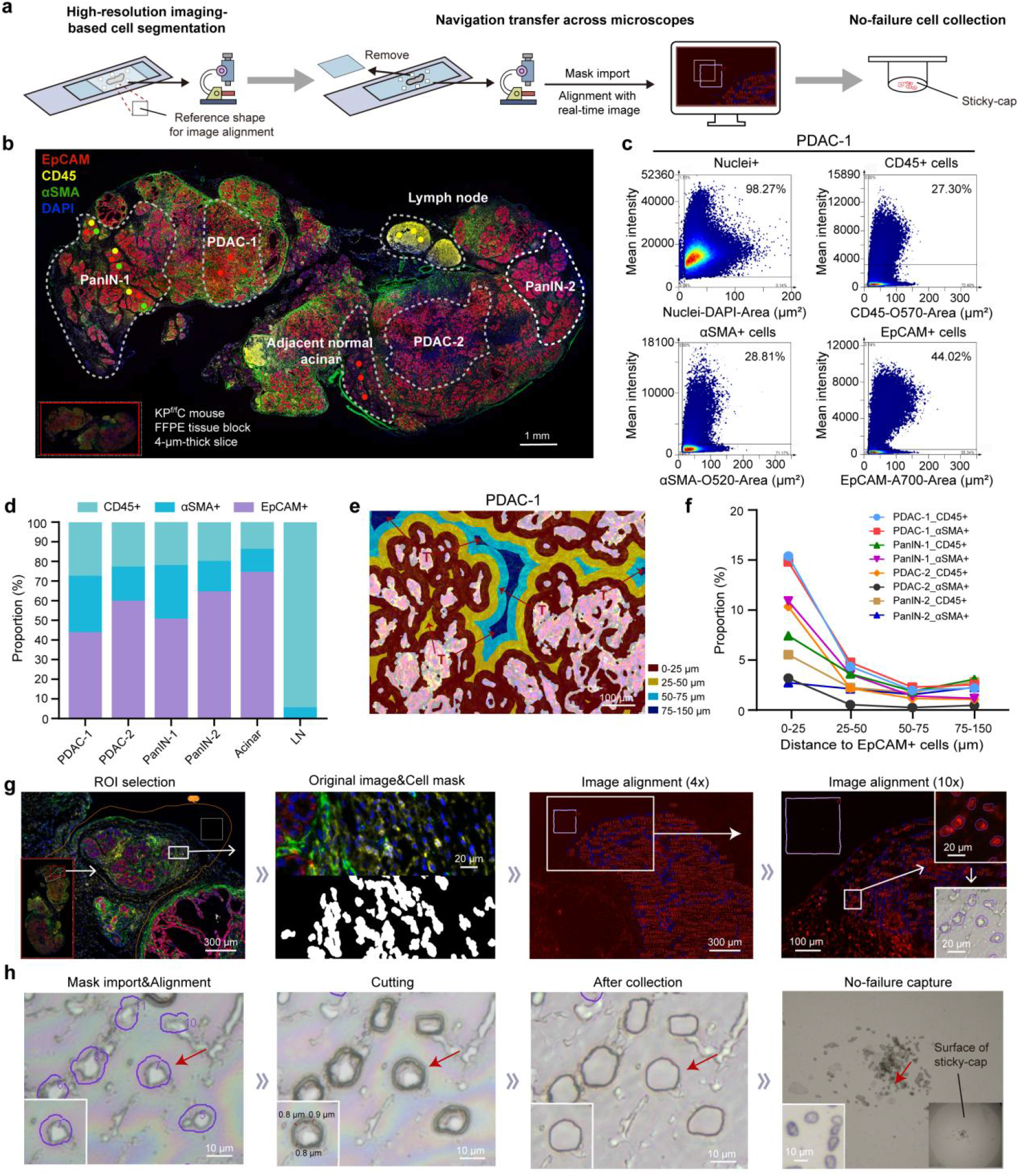
mIHC profiling and LMD capture of single cells in the pancreatic TME. **a**, Spatial proteomics workflow of the SCPro. **b**, Multiplexed immunohistochemical whole-slide image of a 4-μm-thick KP^f/f^C mouse tissue section. The color dot showing the representative LMD cutting position for biological replicates of distinct cell types (n = 3). **c**, Representative tissue cytometry charts of the PDAC-1 region. **d**, Proportion of CD45^+^ immune cells, αSMA^+^ fibroblasts, and EpCAM^+^ epithelial cells in distinct regions of the pancreatic TME. **e**, Distance map showing the spatial distribution of CD45^+^ immune cells and αSMA^+^ fibroblasts with EpCAM^+^ epithelial cells as the center; different distances from tumor cells: 0-25 μm (red); 25-50 μm (yellow); 50-75 μm (light blue); 75-150 μm (dark blue). **f**, Line chart showing the proportion of CD45^+^ immune cells and αSMA^+^ fibroblasts with EpCAM^+^ epithelial cells as the center; different distances from tumor cells. **g**, Workflow of the ROI selection, cell typing and cell mask generation, and image alignment based on the real-time image of LMD. **h**, Automated LMD with single-cell resolution. EpCAM (epithelial cells); CD45 (immune cells); αSMA (fibroblasts); DAPI (nuclei). PanIN, pancreatic intraepithelial neoplasm; PDAC, pancreatic ductal adenocarcinoma; IT, CD45^+^ immune cells in the tumor microenvironment; LN, CD45^+^ immune cells in the lymph node; ROI, region of interest.

The centimeter-scale multi-color whole-slide image comprehensively recapitulates the spatial distribution of distinct pathological features and cell types within the pancreatic TME, including the adjacent normal acinar cells, PanIN, and PDAC (EpCAM^+^ cells), as well as the immune cells (CD45^+^ cells) and CAFs (αSMA^+^ cells) (**Fig. 3b** and **Extended Data** Fig. 5). Furthermore, region-specific distribution of fibroblast-enriched (PanIN-1 and PDAC-1) and fibroblast-deficient areas (PanIN-2 and PDAC-2) were also observed in both PanIN and PDAC stages. Notably, the CD45^+^ immune cells also exhibited a distinct spatial distribution within the TME (IT) and peritumoral lymph nodes (LN). Subsequently, 6 typical regions demonstrating spatial distribution heterogeneity in the pancreatic TME were obtained based on the expression level of surface markers and cellular morphology (**Fig. 3b** and **Extended Data** Fig. 5).

Next, we conducted quantitative tissue cytometry analysis of the 6 cell types in-situ with single-cell resolution (**Fig. 3c** and **Extended Data** Fig. 6). The results showed an increasing proportion of CD45^+^ immune cells and αSMA^+^ fibroblasts surrounding EpCAM^+^ tumor cells during tumor progression from acinar cells to PanIN and finally to PDAC (**Fig. 3d**). Additionally, the PanIN-1 and PDAC-1 regions exhibited a higher proportion of infiltrating immune cells and fibroblasts wrapping compared to the PanIN-2 and PDAC-2 regions, illustrating the distinct spatial distribution of different cell types. The spatial distribution of αSMA^+^ CAFs and CD45^+^ immune cells surrounding the tumor cells is a crucial factor in the prognosis of PDAC^28^. Distance map analysis illustrated a decreasing proportion of CD45^+^ immune cells and αSMA^+^ CAFs as the distance from the tumor increased (**Fig. 3e,f** and **Extended Data** Fig. 7), indicating the formation of stromal barriers by the aggregation of CAFs and suppressive immune cells, particularly in the advanced stages for contributing to the poor prognosis of pancreatic cancer^28^.

After quantitative analysis of the multi-color image in situ, we proceeded to address the next challenge of accurately defining single-cell boundaries and transferring cell typing results for precise automated LMD. Specifically, the boundaries of each cell type were delineated by the StrataQuest (SQ) software^32^ in two steps. Firstly, the nucleus identification algorithm was employed to determine the location of nuclei based on the morphology of nucleus and DAPI staining. Then the cell morphology and staining intensity of surface markers were used to determine the cell boundaries using the membrane identification algorithm (**Methods**). To avoid the damage of cell membrane by the laser of LMD, offset the cell contours for about 1 μm was configured. Finally, a filled mask was built over the corresponding cell types (**Fig. 3g** and **Extended Data** Fig. 8). Subsequently, high-purity cell contours with single cell resolution were generated over the original image (**Fig. 3g**). The obtained cell contours were imported into the LMD and aligned with the real-time image of the LMD microscope using the reference shape generated at the high-resolution imaging stage, by simply “drag and drop” of the shape to the reference square shape in the fluorescence model (**Fig. 3g**). Importantly, the well-annotated single cells were automatically dissected and then collected in real time onto the sticky-cap depressed onto the membrane slide with no-failure under brightfield visualization (**Fig. 3h**).

### SCPro explores spatial proteome heterogeneity of the pancreatic TME

To achieve a balance between sufficient protein identification depth and minimal tissue usage, we isolated 60-100 phenotype-matched cell contours for quantitative proteomic analysis. Benefit from the high sensitivity of the iPAC technology, more than 3,000 proteins and nearly 5,000 proteins were quantified from only 60 cells and 100 cells, respectively (**Fig. 4a**). We first conducted spatial proteomic analysis of three cell types (Acinar cell, PanIN, and PDAC) to study the progression of pancreatic cancer. PCA analysis successfully separated these three neighboring cell types, indicating distinct protein expression profiles (**Fig. 4b**). Differential expression analysis showed more upregulated proteins in acinar cell than PanIN and PDAC regions (**Fig. 4c**), highlighting the heterogeneity between acinar cells and the neoplasia cells, which in line with the fact that acinar cells are normal exocrine cells and the other two cell types are neoplasia cells that exhibit varying degrees of progression^30,33^. GO analysis revealed the enrichment of tumor-related pathways of wound healing and mitochondrial translation in PDAC, and the digestion and pancreatic juice secretion pathways were enriched in acinar cells, which well reflecting their biological functions (**Fig. 4d**)^33,34^. Our dataset also revealed many cell-type specific markers from the proteome of acinar cells, which is consistent with previous studies on the human pancreas (**Fig. 4c**)^33,35^. For instance, Cpa1, a known acinar cell marker playing an importation role in digestive function, exhibited high expression in acinar cell compared to the other two cell types. Additionally, Reg3g, which promotes pancreatic inflammation and tumor progression from acinar cells to PanIN, showed high expression in acinar cells, implying a poor prognosis for acinar cells adjacent to tumor cells (**Fig. 4c** and **Extended Data** Fig. 9)^33,35^.

**Fig. 4.**
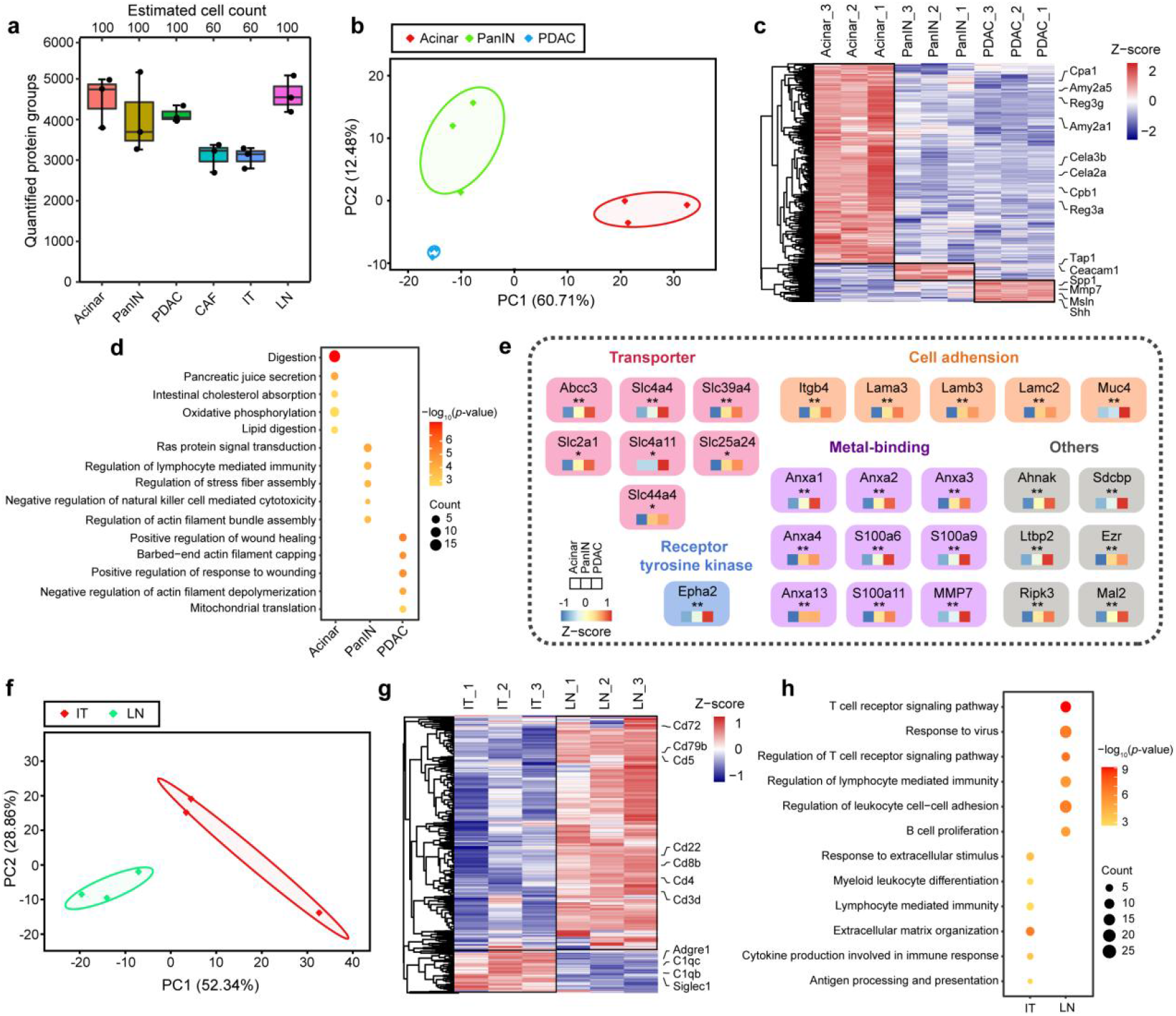
Uncovering the spatial proteomic heterogeneity of the PDAC TME. **a**, Quantified protein groups of each cell type (n = 3). Total area of 120,000 μm^3^ cell contours (∼60 cells) for the CAF and IT regions, and 200,000 μm^3^ cell contours (∼100 cells) for the Acinar, PanIN, PDAC, and LN regions were dissected from the KP^f/f^C mouse tissue section with 4-μm-thick (n = 3). **b**, PCA analysis of the three subtypes of EpCAM^+^ cells. **c**, Heat map showing the significantly differentially expressed proteins for three subtypes of EpCAM^+^ cells (ANOVA, p-value < 0.05, Fold change > 1.2). Protein expression levels were Z-scored. **d**, Dot plot showing the enriched GO biological process terms of the three cell lineages with significantly differentially expressed proteins. **e**, Prognostic markers for pancreatic cancer identified in the dataset and classified by their functions. *, reported in other cancers as a prognostic marker; **, reported in pancreatic cancer as a prognostic marker. (ANOVA, p-value < 0.05, Fold change > 1.2). **f**, PCA showing the distribution of the IT and LN regions. **g**, Heat map showing the significantly differentially expressed proteins of the IT and LN regions (ANOVA, p-value < 0.05, Fold change > 1.2). Protein expression levels were Z-scored. **h**, Dot plot showing the enriched GO biological process terms of the IT and LN regions with the significantly differentially expressed proteins.

PDAC is a highly malignant solid tumor that is typically diagnosed at advanced stages, underscoring the significance of identifying early detection markers for developing treatment strategies. Excitingly, our spatial proteomics data revealed a progressive increase in the expression level of many proteins during PDAC progression on the same tissue slice (**Fig. 4e**). Slc4a4 and Anxa2 which are involved in transport and metal-binding, respectively, have been found to contribute to progression and metastasis of PDAC and have been identified as poor prognosis markers in previous studies (**Extended Data** Fig. 9)^36,37^. In addition to these two proteins, we identified several others that has previously been recognized as prognostic markers for PDAC or other solid tumors^38–40^. Many of these proteins are located on the plasma membrane and play important biological functions, such as transporters, cell-adhesion molecules, and receptor tyrosine kinases (RTKs), making them as potential therapeutic targets (**Fig. 4e**).

Expectedly, spatial proteomic analysis on CD45^+^ immune cells within the tumor microenvironment (IT) and peritumoral lymph node (LN) also revealed significant differences between these two regions (**Fig. 4f**). Differential expression analysis revealed an enrichment of myeloid cell specific markers in the IT (e.g., Adgre1, C1qb, C1qc, and Siglec1), whereas the LN region exhibited an enrichment of lymphoid cell markers (e.g., Cd3d, Cd4, Cd8b, and Cd79b) (**Fig. 4g**). Further GO analysis of the differentially expressed proteins in the IT and LN showed that the upregulated proteins in the IT were associated with myeloid leukocyte differentiation and antigen processing and presentation pathways, while the LN exhibited an enrichment in T cell- and B cell-associated signaling pathways (**Fig. 4h**). These findings demonstrate the distinct spatial distribution of myeloid and lymphoid cell lineages within the tumor immune microenvironment (IT) and peritumoral lymph node (LN), respectively. However, the cell composition information of immune cell subsets in these two regions is limited and needs to be enhanced to further investigate the immune landscape of the pancreatic TME with distinct spatial locations.

### SCPro decodes the pancreatic immune TME through spatial deconvolution

There are numerous cell types in the pancreatic TME, and many of them are present in low abundance while perform crucial functions (e.g., iCAF, apCAF, and Treg)^41,42^. However, the spatial proteomic aspect of SCPro has a relatively low cell-type resolution due to limited abundance of rare cell types and their functional marker signals on a single mIHC-stained slide. To further enhance the resolution of the SCPro for exploring cell subsets in the spatial proteomics data, we seek to generate cell-type specific proteome expression information from the same tumor. Importantly, we adopted spatial deconvolution algorithm which has been widely used in spatial transcriptomics field to systematically explore and correlate the diverse cell type composition in different tissue locations within the pancreatic TME.

To build a comprehensive reference map for spatial deconvolution, we first conducted flow cytometry-based proteomic analysis to acquire cell-type specific proteome data of the main cell types in the pancreatic TME. Herein, 14 distinct cell types, which consisted of CAFs and 3 subtypes of CAFs (myCAFs, iCAFs, and apCAFs), 9 immune cell subpopulations [B cells, CD4^+^ T cells (T4), Tregs, CD8^+^ T cells (T8), myeloid cells (MYE), dendritic cells (DC), macrophages (MAC), neutrophils (NEU), and monocytes (MO)] and the pancreatic cancer cells (PCCs), were included in our flow cytometry-based proteomic analysis (5 biological replicates, total of 69 samples). We optimized our sorting strategy to ensure successful sorting of all the 14 cell types from the single-cell suspension of one KP^f/f^C mouse tumor. To ensure that all the 14 cell types in a viable state could be successfully sorted from the same tumor sample, only up to 1,000 cells for each individual cell type were sorted for further proteomic profiling. Surprisingly, we successfully collected 1,000 cells for all the cell types, excepted for the typically rare cell type myCAF, iCAFs, and apCAFs for which we only collected a few hundreds of cells as expected (**Fig. 5a** and **Extended Data** Fig. 10a,b).

**Fig. 5.**
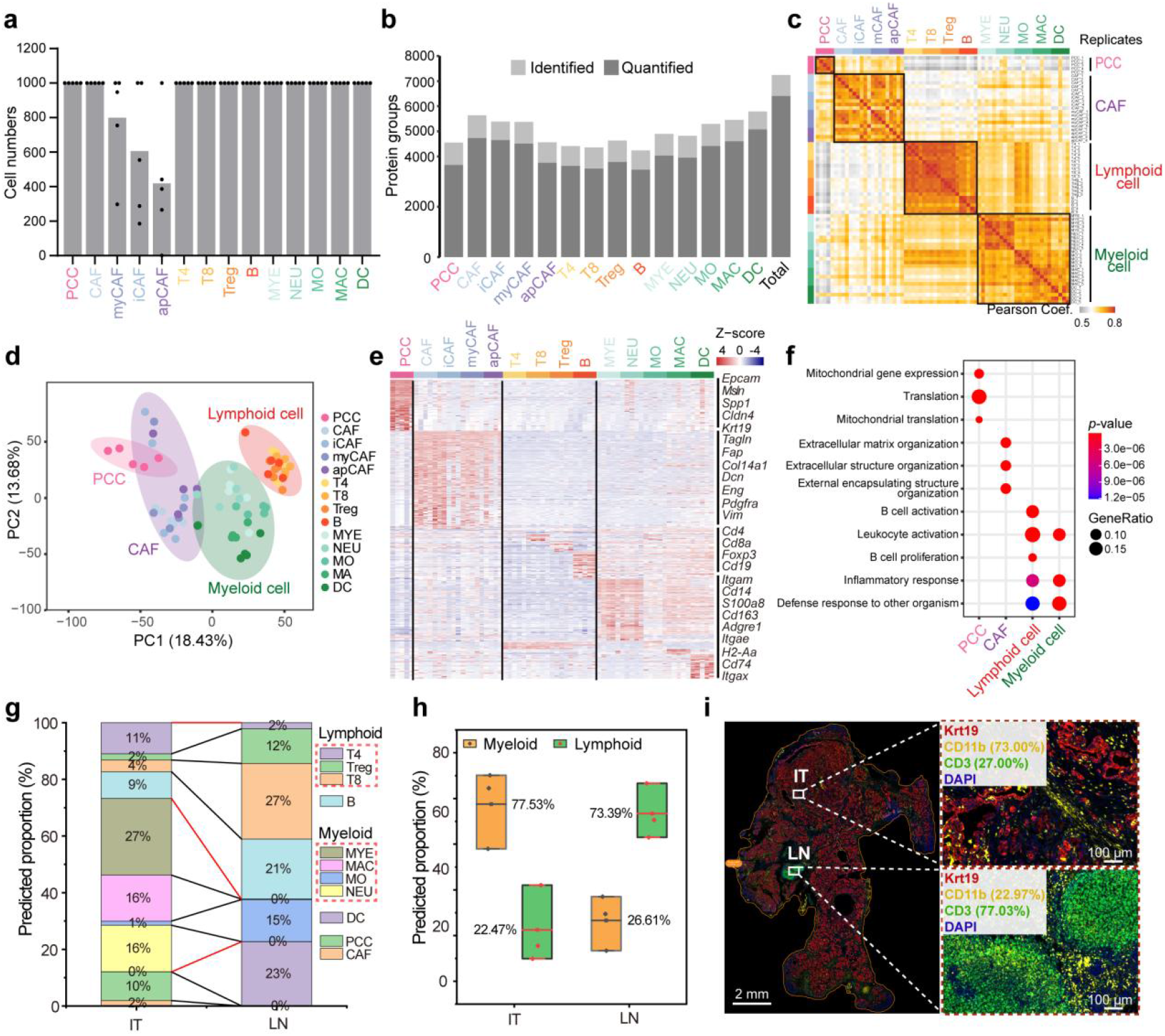
SCPro dissects the PDAC TME through spatial deconvolution. **a**, Cell number of each population acquired by cell sorting. **b**, Proteome coverage for the 14-flow cytometry-sorted cell types, the number of identified and quantified protein groups are indicated in gray and dark, respectively (n = 4 for apCAF and n = 5 for other groups from 5 KP^f/f^C mice). **c**, Pearson’s correlation coefficients of all samples, which are arranged according to their lineage relationships. **d**, PCA plot showing the distribution of measured cell types. Ellipses cover the four cell lineages indicated by colors, involving all cell types. **e**, Heat map showing significantly differentially expressed proteins for each cell type (LIMMA, p-value < 0.05, Fold change > 2). The well-known lineage markers were labeled on the right. Protein expression levels were Z-scored. **f**, Dot plot showing the top enriched GO biological process terms of the 4 cell lineages with significantly differentially expressed proteins. **g**, The predicted proportion of the 14-flow cytometry-sorted immune cell subtypes in the IT and LN regions deconvoluted by Tangram^20^. **h**, Box plot showing the predicted proportion of CD11b^+^ myeloid cells (MYE, MAC, MO, NUE) and CD3^+^ lymphoid cells (T4, Treg, T8) in the IT and LN regions, deconvoluted by Tangram^20^. The sum of the predicted proportions of CD11b^+^ myeloid cells and CD3^+^ lymphoid cells was set to 100% through normalization. **i**, Multiplexed immunohistochemical staining of CD11b^+^ myeloid cells and CD3^+^ lymphoid cells. The sum of the proportion of CD11b^+^ myeloid cells and CD3^+^ lymphoid cells was set to 100% through normalization. Krt19 (epithelial cells); CD11b (myeloid cells); CD3 (T cells); DAPI (nuclei). PCC, pancreatic cancer cell; CAF, cancer-associated fibroblast; iCAF, inflammatory CAF; myCAF, myofibroblastic CAF; apCAF, antigen-presenting CAF; Treg, regulatory T cell; MYE, myeloid cell; NEU, neutrophil; MO, monocyte; MAC, macrophage; DC, dendritic cell.

Leveraging the sensitivity of the iPAC technology, we identified 4,000 to 6,000 protein groups for each cell type and over 7,000 protein groups across all 14 cell types with high reproducibility in ddaPASEF mode (**Fig. 5b,c**). PCA analysis revealed distinct distribution patterns of the proteome among the four lineages (**Fig. 5d**). Lymphoid and myeloid cells showed proximity due to their immune-related characteristics, while non-immune lineages (PCCs and CAFs) exhibited a closer distribution among each other. Notably, apCAFs were closely associated with myeloid cells due to their expression of antigen-presenting proteins^41^. The heatmap of the top differentially expressed proteins for each cell type displayed lineage-specific markers with expected abundance, such as PCCs markers (Epcam, Msln, and Krt19), CAFs markers (Fap, Dcn, and Vim), lymphoid cells markers (Cd4, Cd8, and Cd19), and myeloid cells markers (Itgam, S100a8, and Cd74) (**Fig. 5e** and **Extended Data** Fig. 10c). Further GO analysis of the differentially expressed proteins revealed distinct functions of the four lineages (**Fig. 5f**). For instance, the mitochondrial translation pathway was found to be enriched in cancer cells, indicating its important role in cancer development^34^. Fibroblasts exhibited enrichment in extracellular matrix organization and extracellular structure organization, which well aligns with their critical role in tumor progression through the production of ECM, growth factors, and chemokines^43^. These results well demonstrate the robustness and usefulness of the cell-type proteomics data.

To gain deeper insights into the cellular composition and proportion of CD45^+^ immune cells in the spatial proteome data, we went on to utilize the deconvolution algorithm Tangram^20^ to further decode the proportion of all the 14-flow cytometry-sorted cell subsets in the IT and LN region (**Fig. 5g**). To validate the accuracy of the cell proportion obtained from Tangram, we compared the relative abundance of myeloid cells and lymphoid cells in the IT and LN regions of the pancreatic TME with the cell composition acquired through mIHC imaging by co-staining Krt19 (cancer cells), CD11b (myeloid cells), CD3 (lymphoid cells), and DAPI (**Fig. 5h,i** and **Extended Data** Fig. 11). Notably, the predicted myeloid cells and lymphoid cells proportion is 77.53% and 22.47% in the IT region and 26.61% and 73.39% in the LN region, respectively, which is generally in accordance with the image results, indicating the reliability of the deconvolution algorithm in processing proteome data. It should be noted that our previous bioinformatic analysis of spatial proteomics data also indicates the enrichment of myeloid cell and lymphoid cell lineages in the IT and LN, respectively (**Fig. 4g,h**). In addition, the spatial deconvolution analysis also showed that myeloid cells, including MYE, NEU, and MAC, were the most abundant subtypes of CD45^+^ immune cells in the TME, along with a low fraction of CD8^+^ T cells and DC cells (**Fig. 5g**). This observation further underscored the immunosuppressive nature of the pancreatic TME, which is mainly surrounded by the suppressive myeloid cells^44,45^.

### SCPro enables the discovery of novel cell subtypes in the pancreatic TME

Although spatial and cell-type resolution of the SCPro is greatly improved by integrating the cell-type information based on the proteome information of previously identified cell subsets, such type of cell composition and proportion predication may limit the discovery of biologically novel cell types. To this end, we went on to discover novel sub-cell types in the pancreatic TME by analyzing the plasma membrane (PM) proteins in the cell-type proteomic data. PM proteins play vital roles in tumor ecosystems, which commonly serve as surface markers for distinguishing distinct cell types and represent as valuable therapeutic targets^46^.

We developed a novel bioinformatic strategy to explore reliable surface markers for the identification of novel sub-cell types within the tissue context. The workflow involved in two key procedures (**Fig. 6a**): (1) scoring and ranking cell-type-specific PM proteins by fold change and copy number within the measured cell types; (2) categorizing those ranked proteins according to major biological functions. In addition, we curated a mouse PM proteins database by incorporating Uniprot, Phobius^47^, and DeepTMHMM^48^. The results showed that close 10% of the identified proteins were annotated as PM proteins, and up to 300 PM proteins were identified for most cell populations (**Fig. 6b**). To enhance the differentiation of cell types with functional proximity within the same lineage, we employed two strategies: (1) comparing the measured cell lineages (PCC, CAF, lymphoid cell, and myeloid cell) versus the rest; (2) evaluating individual cell types versus the other cell types within the same lineage. This led to the identification of a significant differential surface marker panel, consisting 206 unique PM proteins across 14 cell types (**Fig. 6c** and **Extended Data** Fig. 12a).

**Fig. 6.**
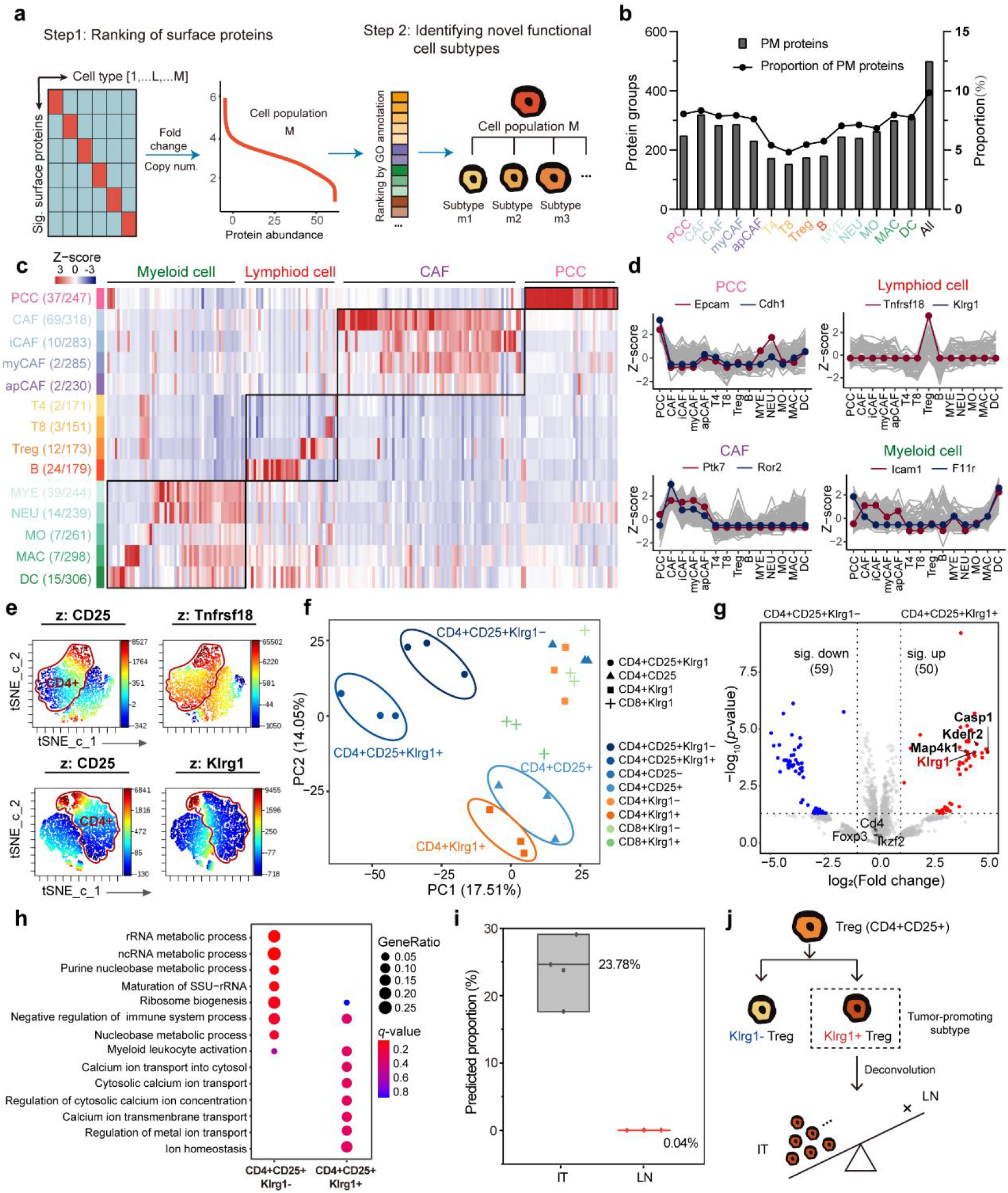
Novel sub-cell type discovery by the SCPro. **a**, Workflow for discovering novel functional sub-cell types from flow cytometry-based proteome data. **b**, Bar plot showing the number of PM proteins. The line chart showing their proportion in all identified protein groups for individual cell types. **c**, Heat map showing the significantly differentially expressed PM proteins (LIMMA, p-value < 0.05, Fold change > 2). Raw names represent the cell type. The number on the left and right of brackets represent the number of significantly differentially expressed PM proteins and total number of PM proteins for each cell type, respectively, ordered by their lineage relationship. Protein expression levels were Z-scored. **d**, Line plot showing the scaled expression levels of significant proteins within four representative cell types (PCC, CAF, Treg, and DC), top 2 of which are colored, respectively. **e**, t-SNE plot showing the expression patterns of Tnfrsf18 and Klrg1 on CD25^+^ Treg cells. **f**, PCA analysis of the 8 Klrg1-associated cell types (n = 3 per cell type from 3 KP^f/f^C mice). **g**, Proteome comparison of CD4^+^CD25^+^Klrg1^-^ Treg (Klrg1^-^ Treg) and CD4^+^CD25^+^Klrg1^+^ Treg (Klrg1^+^ Treg). The significant proteins were showed in blue and red color, respectively (p-value < 0.05, Fold change > 2). **h**, Dot plot showing the enriched GO biological process terms of the CD4^+^CD25^+^Klrg1^-^ Treg (Klrg1^-^ Treg) and CD4^+^CD25^+^Klrg1^+^ Treg (Klrg1^+^ Treg). **i**, Predicted proportion of CD4^+^CD25^+^Klrg1^+^ Treg (Klrg1^+^ Treg) among the 8 Klrg1-associated cell types in the tumor microenvironment (IT) and peritumoral lymph node (LN), respectively, deconvoluted by Tangram^20^. **j**, Summary of discovering a novel Treg subtype and predicting its spatial location.

Based on normalized scores and major biological functions, we highlighted the top 2 differentially expressed PM proteins representing cell types in the four lineages, including known lineage makers and potentially new surface markers for identifying novel sub-cell types (**Fig. 6d**). Interestingly, we found that Tnfrsf18 and Klrg1 were significantly over-expressed on Treg, which is a subset of CD4^+^ T cells with tumor-promoting features and has been linked to a poor prognosis for cancer therapy^49^. The presence of essential functions in Treg was rationalized, as Tnfrsf18 and Klrg1 both function as immune checkpoint molecules^50,51^. To validate our proteomic data, we conducted flow cytometry analysis and verified the elevated expression levels of Tnfrsf18 and Klrg1 on CD25^+^ Treg (**Fig. 6e**). Notably, the t-distributed stochastic neighbor embedding (t-SNE) plot from the flow cytometry analysis showed that CD25^+^ Treg can be subdivided into two subtypes based on the expression of Klrg1 (**Fig. 6e**).

To systematically investigate the role of Klrg1 on T cells, we further sorted 8 of the Klrg1-associated T cell subtypes (i.e., CD4^+^CD25^+^Klrg1^±^, CD4^+^CD25^±^, CD4^+^Klrg1^±^, and CD8^+^Klrg1^±^ T cells) for further cell-type proteomic analysis (**Extended Data** Fig. 12b,c). We then investigated the proteome differences between CD4^+^CD25^+^Klrg1^+^ Treg (also named Klrg1^+^ Treg) and CD4^+^CD25^+^Klrg1^-^ Treg (also named Klrg1^-^ Treg). These two cell types showed a clear distinction in the PCA plot (**Fig. 6f**). The significantly upregulated proteins in the Klrg1^+^ Treg reveals its immunosuppressive and tumor-promoting features (**Fig. 6g**). For instance, Casp1 plays a crucial role in the process of pyroptosis, which is an inflammatory form of cell death^52^. Kdelr2, localized to the ER-Golgi pathway, is associated with the poor prognosis and tumorigenesis of many cancers^53^. Map4k1 (also known as Hpk1) acts as an immunosuppressive regulatory kinase to inhibit the function of T cells and DC, resulting in poorer survival outcomes in pancreatic cancer^54^. Additionally, GO biological analysis also indicated that the Klrg1^+^ Treg plays a more significant role in the activation of myeloid leukocytes (**Fig. 6h**). Last but not the least, to predict the spatial location of Klrg1^+^ Treg, spatial deconvolution was performed utilizing the proteome data of the 8 Klrg1-associated T cell subtypes. The results indicated that the Klrg1^+^ Treg cells were mainly enriched in the IT region rather than the LN region (**Fig. 6i,j**), which correlates well with an increase in the recruitment of myeloid cells within the TME (**Fig. 5h**) and indicates the myeloid cell activation properties of Klrg1^+^ Treg. Collectively, these findings well validate the potential immunosuppressive features of Klrg1^+^ Treg and powerfulness of the multimodal spatial proteomic investigation.

## Discussion

Single cell-resolved spatial proteomic analysis relies on nanoscale processing of limited stained FFPE tissue slice samples. The iPAC technology ensured the sensitivity and versatility of the SCPro platform. Recently, many integrated proteomics sample preparation methods have been proposed to match the increasing sensitivity of advanced MS instruments^1,22^. Based on minimized in-solution digestion concept, most of these methods showed excellent performance in processing flow cytometry-sorted single cells by integrating all sample preparation steps into one pot to minimize sample loss. MS-compatible reagents were used in these methods to avoid interfering MS signal, as they lack subsequent clean-up steps. However, these design seldom fully considered the removal of chemical dyes and other non-protein containments from stained tissue samples, which are the natural components of this type of samples and the major difference with flow cytometry-sorted single cells. The SPE-based iPAC technology addressed these challenges by incorporating the extended wash after protein capture, resulting in enhanced protein coverage for limited stained tissue samples (**Fig. 2d,h**). Our results indicated the cleanness of peptides is equally important to its recovery rate (**Fig. 2e**). Notably, the iPAC technology also showed excellent performance in processing low number of mIHC-stained cells which are LMD-dissected from FFPE tissue section (**Fig. 4a**). Therefore, the iPAC technology is promised to meet the growing demands of spatial proteomic research.

The integration of antibody-guided cell typing technologies and deep proteome profiling, as demonstrated by the SCPro platform, comprehensively uncovers the tissue proteome heterogeneity. The essential advances of the SCPro platform mainly include: (1) the development of an end-to-end workflow for multi-color imaging-based cell segmentation, which enables accurate spatial proteome profiling of multiple cell types on a single FFPE tissue section; (2) the formation of reference shapes for aligning cell contours which enables navigation transfer between high-quality multiplexed imaging with coverslip and low-quality real-time imaging of LMD after removing the coverslip; (3) the dissected cell contours were collected by the sticky-cap directly depressed on the membrane slide under visualization with no-failure, rather than collected by laser pulse or gravity in other LMD systems that may cause sample loss, especially when collecting cell contours with single-cell resolution.

The spatial proteome organization of the pancreatic TME has been studied previously^15,55^, but remained in low cell-type resolution. Guided by the 4-color mIHC image of pancreatic tumor, the SCPro systematically revealed the spatial proteome changes of neoplasia cells and immune cells in the pancreatic TME. Notably, the spatial deconvolution analysis greatly extended the cell-type resolution of the SCPro. For the first time, this study introduced the concept of utilizing flow cytometry-based cell-type resolved proteomics data to infer a more refined cell composition in distinct tissue locations, providing valuable insights into the immune landscape of the pancreatic TME with distinct spatial distribution (**Fig. 5g**). Importantly, by seamless combining the spatial deconvolution analysis with bioinformatic strategy for surface marker screening, we identified a novel sub-cell type of Treg. The spatial location of the novel cell type was predicted to be located in near tumor rather than in the peritumor lymph node (**Fig. 6i**). This finding is insightful, as the previous report concluded that the completely depletion of Treg accelerated tumorigenesis due to compensatory myeloid infiltration^41^. Our results suggested the more aggressive subtype of Treg found in pancreatic TME as a potential therapeutic target.

Collectively, our study presents a streamlined proteomic workflow for advancing our understanding of tissue heterogeneity in spatial context. The SCPro extends the comprehensiveness of traditional digital histopathology by incorporating the proteomic dimension. Along with future advancements of LC-MS instrumentation and data mining algorithms, the SCPro is expected to become a generic tool for systematic characterization of spatiotemporal proteomic landscape and cell-cell interaction within TME at nanometer resolution.

## Methods

### Animal experiment and ethics

Mouse brains were obtained from 6- to 8-week-old C57BL/6 mice. Detail protocols for the attachment of mouse brain sections and H&E staining were described previously^55^. Kras^LSL-G12D/+^; Trp53^flox^; Pdx1-Cre mice (denoted as KP^f/f^C mice) were bred and raised in the Animal Experiments Center at Southern University of Science and Technology. The genotyping of KP^f/f^C mice was determined by PCR using tail biopsies as described previously^29^. To harvest the KP^f/f^C tumors, the KP^f/f^C mice were sacrificed by cervical dislocation, the tumors were then removed, washed with ice-cold phosphate buffered saline (PBS) for three times, transferred to a 4% paraformaldehyde solution for fixing for 24 to 48 hours. After fixation, the tissues were paraffin-embedded for further analysis. All protocols regarding mouse experiments were approved by the Institutional Animal Care and Use Committee at Southern University of Science and Technology of China.

### Multiplexed immunohistochemical staining

A 4-μm-thick tissue sections were cut using a microtome (Leica) and mounted onto the frame slides. To ensure optimal staining, the frame slides underwent a deparaffinization process by incubating for 10 minutes in 100% xylene for three times, and rehydrated by a series of 100%, 100%, 90%, 80%, and 70% ethanol, each for 5 minutes, then the tissue sections were washed with ddH_2_O for 5 minutes. The mIHC staining was conducted using the TSA kit (TissueGnostics, TGFP7100) following the manufacturer’s instructions. Primary antibodies, Anti-EpCAM (Cell Signaling Technology, clone E6V8Y, dilution 1:500), anti-CD45 (Cell Signaling Technology, clone D3F8Q, dilution 1:1000), anti-αSMA (Cell Signaling Technology, clone D4K9N, dilution 1:500), anti-Krt19 (Cell Signaling Technology, clone D4G2, dilution 1:1000), anti-CD3e (Cell Signaling Technology, clone D4V8L, dilution 1:100), and anti-CD11b (Cell Signaling Technology, clone E6E1M, dilution 1:1000) were used for staining KP^f/f^C mouse tumor sections. Finally, the samples were mounted with DAPI Fluoromount-G® antifade mountant (SouthernBiotech, 0100-20) and coverslips to obtain high-quality images for further analysis.

### Multiplexed immunohistochemical image acquisition and analysis

Before image acquisition, the square reference shapes for image alignment were marked onto the membrane slide by LMD. The whole-slide image was first acquired by the TissueFAXS Spectra Systems (TissueGnostics) using a 5× objective to identify the location of the tissue section. Then the multiplexed whole-slide images were acquired at 40× high-magnification, and greyscale images of high magnification were extracted for each dye channel for further analysis. The StrataQuest software version 7.1 (TissueGnostics) was used for the quantitative analysis and cell typing of the high magnification images. For the cell typing of KP^f/f^C tumor tissue section, the nuclei identification algorithm was used to identify the nucleus. The parameter for nuclei size was set at 10 pixels. Then, the cell membrane identification algorithm was utilized to identify of the cell boundaries of corresponding cell types. The parameters for cell membrane identification were set at -0.32 μm interior radius, 0.63 μm exterior radius, and 4 μm maximum growing step. After identifying the nuclei and cell membrane, a filled mask was generated over the original image of the corresponding cell type. Then, the cutting path was offset for about 1 μm. The square reference shapes were then exported with the cell contours after cell typing to guide automated LMD.

### Laser microdissection

The CellCut system (MMI) was used to collect cell contours. The mask files exported from the StrataQuest software were imported into the CellCut system to guided automated LMD. The clearly recognizable square reference shape was used to ensure the alignment of the cell contours generated from the StrataQuest software with the real-time image of the LMD under fluorescence mode. The cell contours were cut at 40× objective in brightfield mode. The cell contours were collected using the IsolationCap (MMI) and stored at -20 °C for further analysis.

### Tumor dissociation and cell sorting

The tumor resected from KP^f/f^C mouse was washed with ice-cold PBS to remove redundant fat and vessels. Then the tumor was minced into 2-4 mm pieces, and transferred into the gentleMACS C tube (Miltenyi, 130-096-335) with the enzyme mix solution from the tumor dissociation kit (Miltenyi, 130-096-730) prepared following the manufacturer’s instructions. The tumor was dissociated using the gentleMACS^TM^ dissociator (Miltenyi). The cell suspension was then filtered through a 70 μm cell strainer (Corning, 431751) and washed twice by ice-cold RPMI-1640 (Corning, 10-040-CMR) to obtain a single-cell suspension. The single-cell suspension was centrifuged at 300 g for 5 minutes at 4 °C and the supernatant was completely aspirated. One milliliter of stain buffer (BD Pharmingen, 554657) was used to resuspend the cell precipitation. The cell number was counted and divided into three panels for cell staining to reduce sample loss. The detailed information about the antibodies used in flow cytometry and the panel design are showed in **Extended Data Tables 1 and 2**. The cells were stained according to the manufacturer’s instructions and sorted using the BD FACSAria SORP flow cytometer (BD Biosciences). The cells were collected into 1.5 mL Protein LoBind tubes (Eppendorf, 022431081) using the 4-way sorting and single-cell mode, at 4 °C. In order to reduce potential cell loss, the cells were centrifuged at 2000 rpm for 5 minutes at 4 °C after sorting, flash freezed down in liquid nitrogen, and stored at -80 °C freezer for further analysis.

### Sample preparation by iPAC

For tissue and sorted cell samples, 20 μL of lysis buffer composed of 1% (w/v) DDM, 10 mM HEPES (pH 7.4), 150 mM NaCl, 600 mM guanidine HCl, and a protease inhibitor mixture (Roche) was used. The collected tissue slices were sonicated in the lysis buffer with the non-contact sonication using Bioruptor (Diagenode) for 20 cycles (30 s-on, 30 s-off) at 4 °C. Two plugs of C18 disks (3M Empore) and two plugs of SAX disks (3M Empore) were inserted into 200 μL pipette tips to fabricate the iPAC spintips. Before sample loading, the tips were equilibrated with DDM coating buffer (0.1% DDM in NH_4_OH) with a brief centrifuge, then the equal volume of tissue or cell lysates was mixed and loaded with the DDM coating buffer. Afterward, protein aggregation on the SAX disks was induced by loading and incubating in pure ACN for 10 minutes. The samples were then subjected to extended wash with 80% (v/v) ACN twice. The proteins were reduced using 50 mM dithiothreitol (DTT) in 20 mM ammonium bicarbonate (ABC) and incubated for 30 minutes at 37 °C. Specifically, 4 μL of digestion buffer containing 20 ng/μL sequencing-grade trypsin (Promega), 20 ng/μL sequencing-grade Lys-C (Wako), and 10 mM iodoacetamide (IAA) in 20 mM ABC were added to the tips and incubated in darkness for 3 hours at 37℃ for digestion. The digested peptides were then transferred to the C18 layer of the iPAC tip through 60 μL 1 M NaCl in 1% (v/v) formic acid (FA). After desalting with 60 μL 1% (v/v) FA for twice, the resulted clean peptides were eluted by 60 μL 80% (v/v) ACN into glass insert. Peptides were lyophilized to dryness for MS analysis.

### High-pH reversed-phase chromatography fractionation

We conducted high-pH reversed-phase chromatography fractionation to generate a deep proteome library of KP^f/f^C mouse tissues for data-independent analysis (DIA). Around 100 μg of peptides from 5 KP^f/f^C mice tumor sections were fractionated on an XBridge peptide BEH C18 column (130 Å, 5 μm, 2.1 mm × 150 mm) using a 60 minute-gradient and concatenated into 24 fractions on an microflow HPLC (Agilent 1260). The peptide samples were vacuum dried in a SpeedVac (Thermo Fischer), then reconstituted in 0.1% FA spiked with iRT peptides (Biognosys) for LC-MS/MS analysis.

### Liquid chromatography

The lyophilized peptides were reconstituted in 2.5 μL of 0.1% (v/v) FA. Only 2 μL of the redissolved peptides were injected for the single-shot LC-MS/MS analysis. A homemade 50 μm I.D. × 20 cm separation column with integrated fritted tip was used by packing with 1.9 μm C18 beads (Dr. Maisch) and coupling to a nanoElute liquid chromatography system (Bruker Daltonics). The temperature of the separation column was maintained at 50 °C using an integrated column oven. Mobile phases A and B consisted of 0.1% FA and ACN, respectively. A segmented 80-minute gradient was used for LC-MS analysis. The gradient was set as follows: from 2 to 22% (v/v) buffer B in 50 minutes, from 22 to 35% (v/v) buffer B in 10 minutes, from 35 to 80% (v/v) buffer B in 10 minutes, holding at 80% (v/v) buffer B for the last 10 minutes.

### Data acquisition in DDA mode and DIA mode

The timsTOF Pro (Bruker Daltonics) was used to analyze the eluted peptides. For DDA acquisition, the scan range was set to m/z 300-1500 in the positive mode. The ramp time was 200 milliseconds, and the total cycle time was 1.03 seconds with one MS scan and 4 parallel accumulation-serial fragmentation (PASEF) scans. The ion mobility (1/K_0_) was scanned from 0.75 to 1.30 Vs/cm^2^. For DIA acquisition, the dia-PASEF method was optimized using the py_diAID software^56^ with the m/z range of 300 to 1500, the ion mobility range was set to 0.75 to 1.30 Vs/cm^2^, the ramp time was 200 milliseconds. Each dia-PASEF scan with variable isolation window widths that were adjusted according to the precursor densities. The optimized method includes two ion mobility windows and 12 dia-PASEF scans^57^.

### Raw data analysis

All the MS raw files for the iPAC benchmark were acquired in DDA mode and defaulted LFQ-MBR workflow of MSFragger software (version 3.7) integrated in Fragpipe platform (version 19.0). The MS raw files were searched against the reviewed mouse UniProt FASTA database (21,984 entries). The spatial proteomic data of KP^f/f^C mice were acquired in DIA mode and searched by Spectronaut (version 17.4). The MS raw files were searched against the reviewed mouse UniProt FASTA database (21,984 entries). All the raw files of flow cytometry-based proteomic data of KP^f/f^C mice were acquired in DDA mode and searched by the MSFragger software (version 3.5) integrated in Fragpipe platform (version 17.0). The MS raw files were searched against the reviewed mouse UniProt FASTA database (17,101 entries). Trypsin and LysC were set as digestion enzymes and a maximum of two missed cleavages were allowed for all the raw files.

All flow cytometry raw data were saved as .fcs files and processed by the FlowJo software (version 10, BD Biosciences) for further analysis. For the t-SNE plot analysis, the .fcs files were upload to the Cytobank (https://www.cytobank.cn/). The t-SNE-CUDA algorithm was used in the dimensionality reduction analysis. An equal number of 50,000 cells were randomly selected for the analysis of each sample. The perplexity parameter was set according to the default factory setting.

### Bioinformatic processing of proteomics data

The contamination ratios of the heatmaps were defined as the ratio between the summed intensity of the up-left and bottom-right precursors in the heatmap. The white lines in the figures pass through points of (350m/z, 0.8IM) and (950m/z, 1.3IM). Up-left precursors were often regarded as +1 contaminants^27^. Precursors were extracted by AlphaTims software (version 1.0.7)^58^. This calculation was performed using a custom R programming script. Furthermore, the total MS intensity at specific retention times and the distribution of PSM were calculated based on the CSV files exported from AlphaTims.

Proteomics data analysis and visualization were performed using Perseus software (2.0.7.0)^59^ and R Studio (4.1.0). After excluding potential contaminants, we filtered the quantified proteins for at least two valid values in at least one cell type. Missing values were imputed using minimum valid values of 0.1. LIMMA package (3.48.0)^60^ and clusterProfiler package (4.0.5)^61^ in R were used for statistical analysis and GO enrichment analysis, respectively.

To construct membrane proteins databases, we first downloaded the reviewed mouse proteins sequence from Uniprot (17,119 entities), and retained 4,487 proteins annotated with “transmembrane”. Then, Phobius^47^ and DeepTMHMM^48^ were individually used to predict transmembrane proteins with previous downloaded protein sequences. Proteins predicted as “transmembrane” by less than two approaches were discarded. Combined with reported surfaceome^62^, our membrane protein databases involved 4,518 high quality proteins, 38% of which were detected in this study.

We performed deconvolution of the raw mouse pancreatic cancer spatial data using the Tangram^20^ (version 1.0.4) on log2-normalized mouse pancreatic cancer cell-type proteomics data on top 1000 proteins, and other parameters were set to default.

## Acknowledgements

The authors acknowledge funding from the China State Key Basic Research Program Grants (2020YFE0202200, 2022YFC3401104, 2021YFA1301601, 2021YFA1301602 and 2021YFA1302603), the National Natural Science Foundation of China (92253304, 22125403, 2201218, and 22104047), the Shenzhen Innovation of Science and Technology Commission (JSGGZD20220822095200001, JCYJ20200109141212325, JCYJ20210324120210029 and JCYJ20200109140814408), and Guangdong province (2019B151502050).

## Author contributions

Y.X. investigation, performed experiments, analyzed data and wrote the paper. X.W. performed experiments, analyzed data and wrote the paper. Y.L. conducted the bioinformatic analysis and wrote the paper. Y.M. performed the experiments, analyzed data, and wrote the paper. Y.S. and Y.Y. experiment design and performed experiments. W.G. provided advice for the data analysis and data organization. C.F. provided necessary reagents and gave advice for the experiment design. W.C. gave advice for the experiment design. X.Y., F.L. and P.B. gave advice for the experiment design and technical support. Y.S. and R.X. gave advice for the experiment design. R.T. conceived and supervised the project, designed the experiment and wrote the paper.

